# Lack of mRNA Methylation in Schwann Cells Results in Demyelination and Regenerative Failure

**DOI:** 10.1101/2025.06.03.657089

**Authors:** Mehmet Can Sari, Anna Johnson, Andrew Tian Yu, Ruifa Mi, Warren Chen, Xindan Hu, Thomas GW Harris, Sami Tuffaha, Vivek Swarup, Riki Kawaguchi, Guo-Li Ming, Ahmet Hoke

**Author notes:** **Correspondence:** Ahmet Hoke, Johns Hopkins School of Medicine, 855 N. Wolfe St., Rangos 248, Baltimore, MD, 21205, USA.

## Abstract

Schwann cells are essential for peripheral nerve myelination and regeneration. N6-methyladenosine (m6A) RNA methylation, regulated by methyltransferase-like 14 (Mettl14), is a critical post-transcriptional modification, but its role in Schwann cell biology remains unclear. Using a conditional knockout (cKO) mouse model, we investigated the impact of Mettl14-mediated m6A methylation on Schwann cells. Mice born with Schwann cell-specific genetic deletion of Mettl14 developed normally but starting in young adulthood exhibited progressive motor deficits, severe demyelination, and axonal degeneration, confirmed by behavioral assessments and histological analyses. Mettl14-deficient Schwann cells displayed impaired proliferation and mitochondrial dysfunction in vitro. Following sciatic nerve injury, Mettl14 cKO mice showed defective macrophage recruitment, slowed axonal degeneration, and impaired regeneration. These findings suggest that Mettl14-mediated m6A methylation is critical for Schwann cell maintenance but not development. Given that Mettl14 cKO mice developed a demyelinating polyneuropathy, it is possible that manipulation of m6A methylation in Schwann cells is a promising therapeutic strategy targeting peripheral nerve repair and myelination.

## Introduction

Schwann cells, highly adaptable glial cells in the peripheral nervous system (PNS), not only form and maintain the myelin sheath but also play a crucial role in peripheral nerve regeneration by initiating Wallerian degeneration, facilitating immune cell recruitment for myelin debris clearance, promoting axonal regrowth through dedifferentiation and secretion of neurotrophic factors. (1,2) Peripheral nerve regeneration is a highly coordinated process, distinguished by its remarkable regenerative capacity compared to the central nervous system (CNS). However, factors such as injury severity, the distance between the injury site and the target tissue, and age can affect the success of regeneration (3,4).

RNA modifications are chemical alterations to RNA molecules that impact their stability, translation, splicing, and overall function. More than 170 distinct RNA modifications have been found, with some of the most common ones being N6-methyladenosine (m6A), pseudouridine (Ψ), and 5-methylcytosine (m5C). These modifications are dynamic and reversible, allowing cells to quickly adapt RNA function in response to environmental and cellular signals (5). Among these, m6A methylation is one of the most prevalent and well-characterized RNA modifications in eukaryotic cells. It plays a crucial role in post-transcriptional regulation of RNA metabolism, influencing RNA stability, splicing, translation, and degradation. The m6A modification is dynamic and reversible, controlled by three key classes of proteins: “writers,” “erasers,” and “readers” (6).

The primary m6A “writer” complex includes methyltransferases such as METTL3 and METTL14, which catalyze the addition of m6A marks to specific RNA transcripts. Other proteins, like WTAP and RBM15, assist in the localization and activity of this complex. On the other hand, “erasers” such as FTO and ALKBH5 remove m6A modifications, making the process reversible. Finally, “readers,” including YTH domain-containing proteins (YTHDF1, YTHDF2, YTHDC1), recognize and bind to m6A-modified RNA, influencing processes such as translation, degradation, and RNA localization (7-11).

m6A methylation plays a crucial role in various biological processes, including neurogenesis, and synaptic plasticity (12). Recent studies in oligodendrocytes have shown that loss of Mettl14 disrupts myelination in the central nervous system (CNS), underscoring the importance of m6A in neuronal development (13). Given the dynamic role of m6A methylation in regulating gene expression and cellular function, its dysregulation is noted in a wide range of diseases, including neurological disorders and cancer (7). Understanding the interplay between m6A “writers,” “erasers,” and “readers” provides key insights into RNA-level gene regulation and highlights potential therapeutic avenues (14).

To investigate the function of m6A in PNS myelinating cells, we employed a conditional knockout (cKO) mouse model for Mettl14. Mice carrying a conditional allele of Mettl14 (Mettl14^flox/flox^) were crossed with mice expressing Cre recombinase under the myelin protein zero (P0) promoter, enabling us to specifically investigate m6A’s role in Schwann cell development and function. We performed behavioral and histological assessments in mice, and Schwann cell proliferation and mitochondrial activity in vitro. We conducted sciatic nerve injury experiments to assess degeneration, regeneration, and macrophage recruitment, while bulk RNA sequencing identified key genes and transcription factors involved in these processes.

Our findings suggest that mRNA methylation doesn’t appear to play a role in Schwann cell development but plays a crucial role in Schwann cell maintenance and peripheral nerve regeneration. While Mettl14 cKO mice exhibit normal initial development, they progressively develop demyelination, regenerative failure, and Schwann cell proliferation defects, highlighting the significance of m6A in peripheral nerve function.

## Results

### Mettl14 cKO mice develop progressive neuropathy

To investigate the role of m6A methylation in Schwann cell development and function, we utilized a P0-Cre x Mettl14 conditional knockout (cKO) mouse model. Behavioral assays were conducted monthly from 3–4 weeks to 12 months of age on wild-type (WT, C57BL/6J) and Mettl14 cKO mice to evaluate neuromuscular and motor functions. These assessments included neuromuscular SHIRPA (NM-SHIRPA) scoring for overall neuromuscular function, the accelerating rotarod test for motor coordination and balance, and grip strength measurements of forelimbs and hindlimbs to evaluate skeletal muscle strength.

NM SHIRPA scoring revealed no neuromuscular deficits in Mettl14 cKO mice up to 3 months of age. However, progressive dysfunction emerged thereafter, with NM-SHIRPA scores significantly elevated in Mettl14 cKO mice compared to WT controls starting at 4 months (p < 0.001, n = 7; Figure 1A). Motor coordination and balance, assessed through the accelerating rotarod assay, showed a significant reduction in latency to fall in Mettl14 cKO mice as early as 3 months (p < 0.0001, n=9), whereas WT controls maintained stable performance throughout the study (Figure 1B).

**Figure 1.**
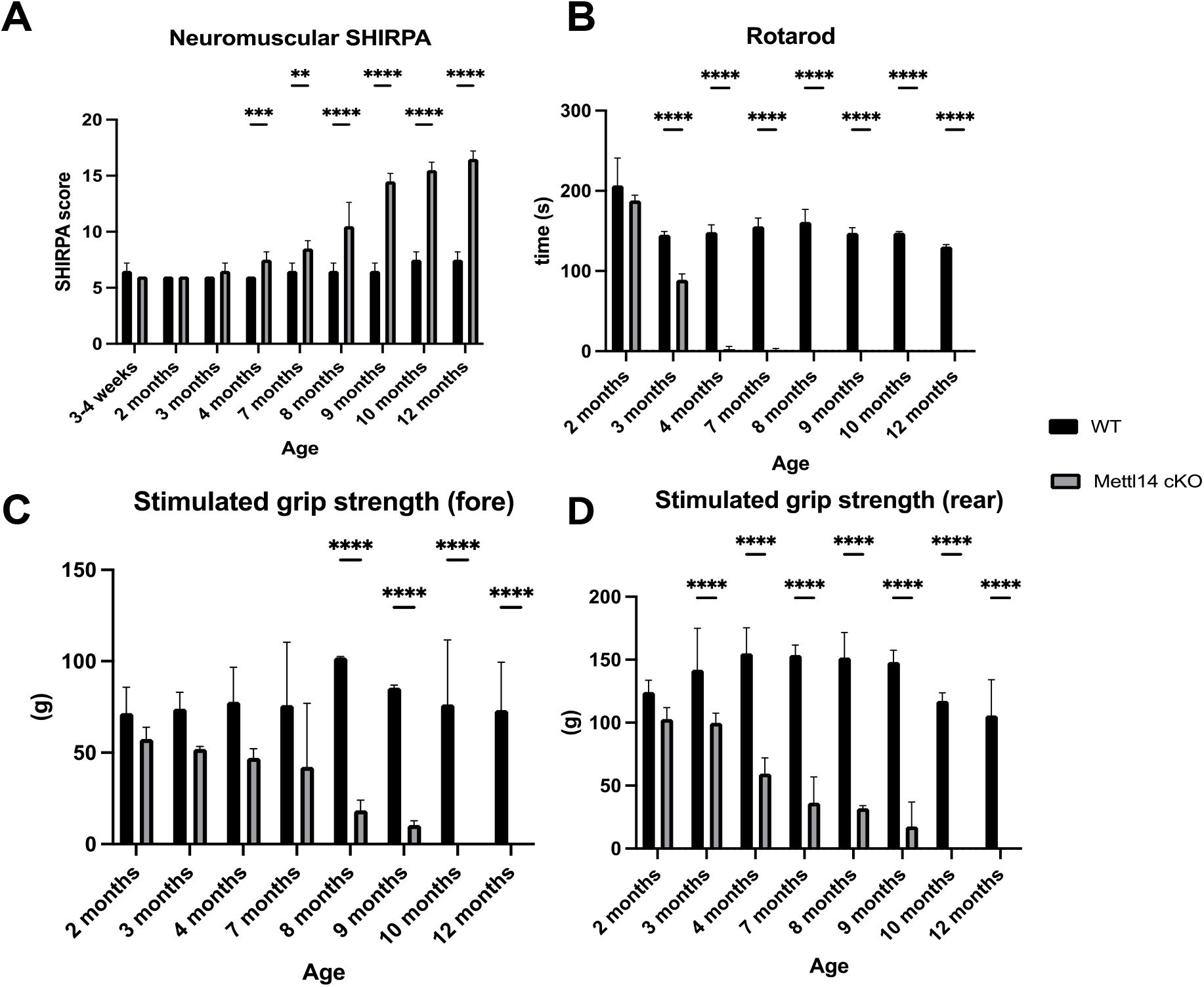
Clinical characterization of mice. Mettl14 conditional knockout mice do not demonstrate behavioral deficits by 4 months of age. 3-4 weeks to 12 month-old wild type and Mettl14 cKO mice were evaluated by (A) Neuromuscular SHIRPA, (B) accelerating rotarod assay, (C) forelimb grip strength test, (D) rearlimb grip strength test. n= 3-20 animals per each time point group, two-way ANOVA, Sidak test, ****p < 0.0001 ; ***p < 0.001 ; **p < 0.01 : ns (non significant) p > 0.05. WT,wild type; Mettl14 cKO, Mettl14 conditional knock out.

Forelimb grip strength remained comparable to WT controls until 7 months, after which it significantly declined in Mettl14 cKO mice (p < 0.0001, n = 5). In contrast, hindlimb grip strength deficits were evident as early as 3 months (p < 0.0001, n = 9; Figures 1C, 1D). Both forelimb and hindlimb grip strength were comparable to WT controls prior to these respective time points. Gender-specific analyses revealed no significant differences in any measured parameters between male and female mice (Supplemental Figures S1, S2).

These findings indicate that Mettl14 deletion in Schwann cells does not appear to impair early development but leads to progressive neuromuscular and behavioral deficits starting at 3-4 months. This underscores the critical role of m6A methylation in Schwann cell maintenance and function beyond early developmental stages.

To investigate the effects of Mettl14 deletion on Schwann cells and peripheral nerve morphology, we performed electron microscopy on sciatic nerves from WT and Mettl14 cKO mice at 21 days, 4 months, and 12 months—key stages representing peripheral nerve myelination and aging. At 21 days, a developmental milestone marking the completion of myelination, both WT and Mettl14 cKO mice exhibited normal sciatic nerve morphology. Quantitative and qualitative analyses showed no significant differences in myelin thickness or axon density between the two groups (Figures 2A, B, C, D, E). However, by 4 months, histological abnormalities started to emerge in Mettl14 cKO mice, coinciding with mild clinical symptoms such as tremors and hindlimb clenching (Movie S1). G-ratio analysis revealed significantly thinner myelin sheaths in Mettl14 cKO mice compared to WT controls (p < 0.001, n = 4), along with reduced axon density and a marked loss of myelinated axons (Figures 2B, C, D).

**Figure 2.**
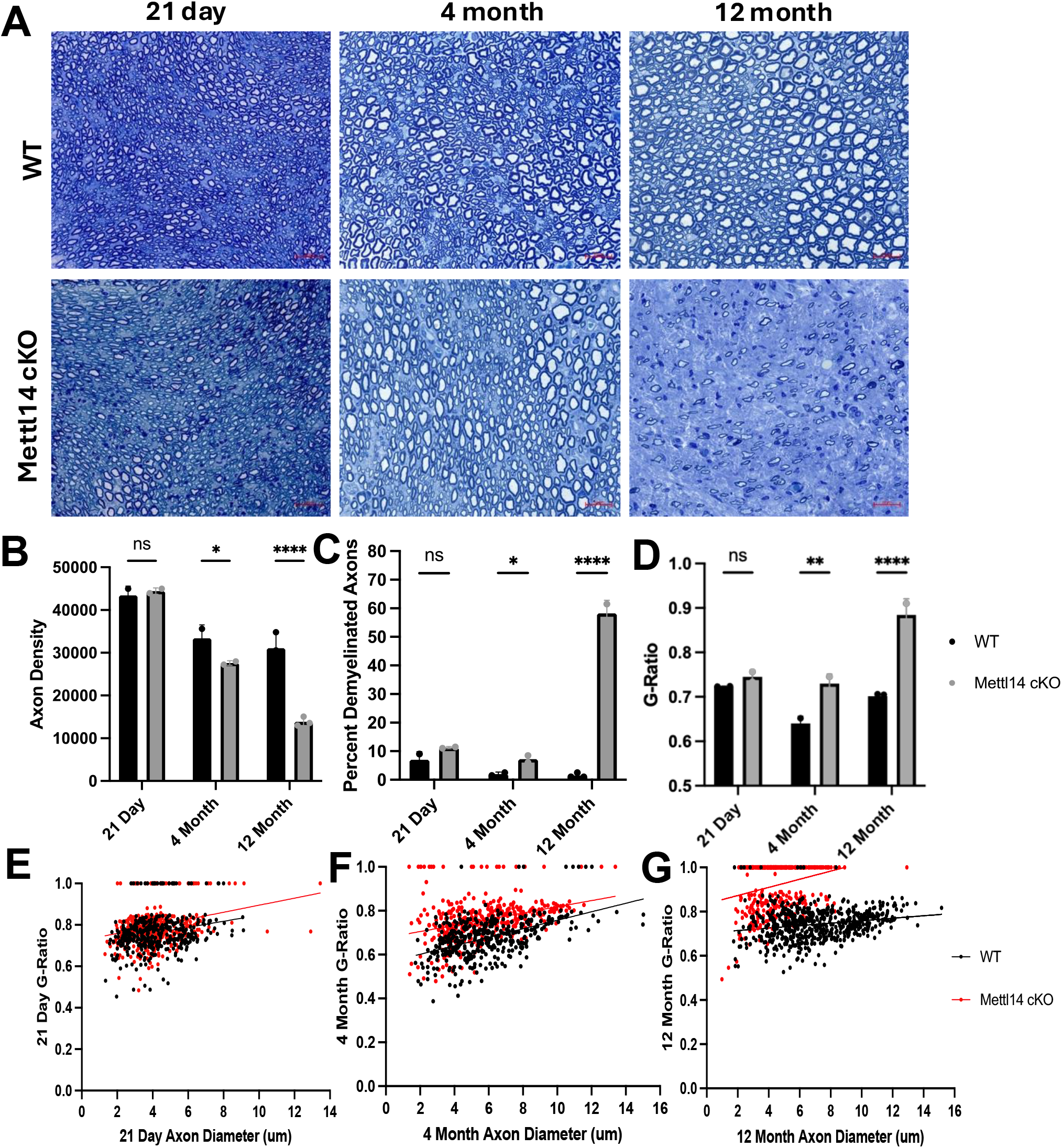
Histological characterization of mice. Sciatic nerves were harvested from 21-day-old, 4-month-old, and 12-month-old wild type and Mettl14 cKO mice and plastic sections were prepared for electron microscopy. (A) Representative images, scale bar 20 μm. Nerve morphometry was performed and axon density (B), percent demyelinated axons (C) average G-ratios (D), and scatter plots G-ratio against axon diameter (E, F, G) are displayed. n= 2-4 animals per condition (>200 axons per animal), two-way ANOVA, simple linear regression, ****p < 0.0001 ; **p < 0.01 ; *p < 0.05 ; ns (non significant) ; p > 0.05. WT,wild type; Mettl14 cKO, Mettl14 conditional knock out.

By 12 months, Mettl14 cKO mice exhibited severe tremors and ataxia (Movie S1), consistent with advanced neuropathological changes. Electron microscopy revealed extensive demyelination, a significant reduction in the number of myelinated axons, and an elevated g-ratio (0.85, p < 0.0001, n=3), reflecting severe myelin thinning (Figures 2C, D). Scatter plots of G-ratio versus axon diameter confirmed a disproportionate loss of large-caliber axons in Mettl14 cKO mice (Figure 2G). Western blot analysis of sciatic nerve lysates further supported these findings, demonstrating significantly reduced levels of myelin basic protein (MBP) in Mettl14 cKO mice by 4 months (p < 0.0001, n=2 Supplemental Figures S3). Collectively, these findings demonstrate that Schwann cell-specific deletion of Mettl14 disrupts myelin integrity and axonal health, leading to progressive demyelination and axonal loss.

To evaluate the effects of Mettl14 deletion on neuromuscular junction (NMJ) innervation, we analyzed NMJs in the soleus muscle of 6-month-old WT and Mettl14 cKO mice. NMJs were visualized using α-bungarotoxin to label acetylcholine receptors (AChRs), while presynaptic terminals were labeled with SMI312 (neurofilaments in axons) and SV2 (synaptic vesicles). Based on the degree of overlap between presynaptic terminals and postsynaptic AChRs, NMJs were categorized as intact (>95% overlap), partially denervated (5–95%), or fully denervated (<5%). In WT mice, NMJs displayed near-complete overlap between presynaptic terminals and postsynaptic AChRs, indicative of proper synaptic function and stability. In contrast, Mettl14 cKO mice exhibited a significant reduction in intact NMJs (p < 0.01), alongside a marked increase in fully denervated NMJs. Partially denervated NMJs were also significantly more frequent in Mettl14 cKO mice, reflecting progressive synaptic denervation (Figures 3A, B). These findings underscore the essential role of Mettl14 in maintaining NMJ innervation. The observed increase in denervated NMJs in Mettl14 cKO mice likely contributes to impaired neuromuscular transmission and the motor deficits documented in this model.

**Figure 3.**
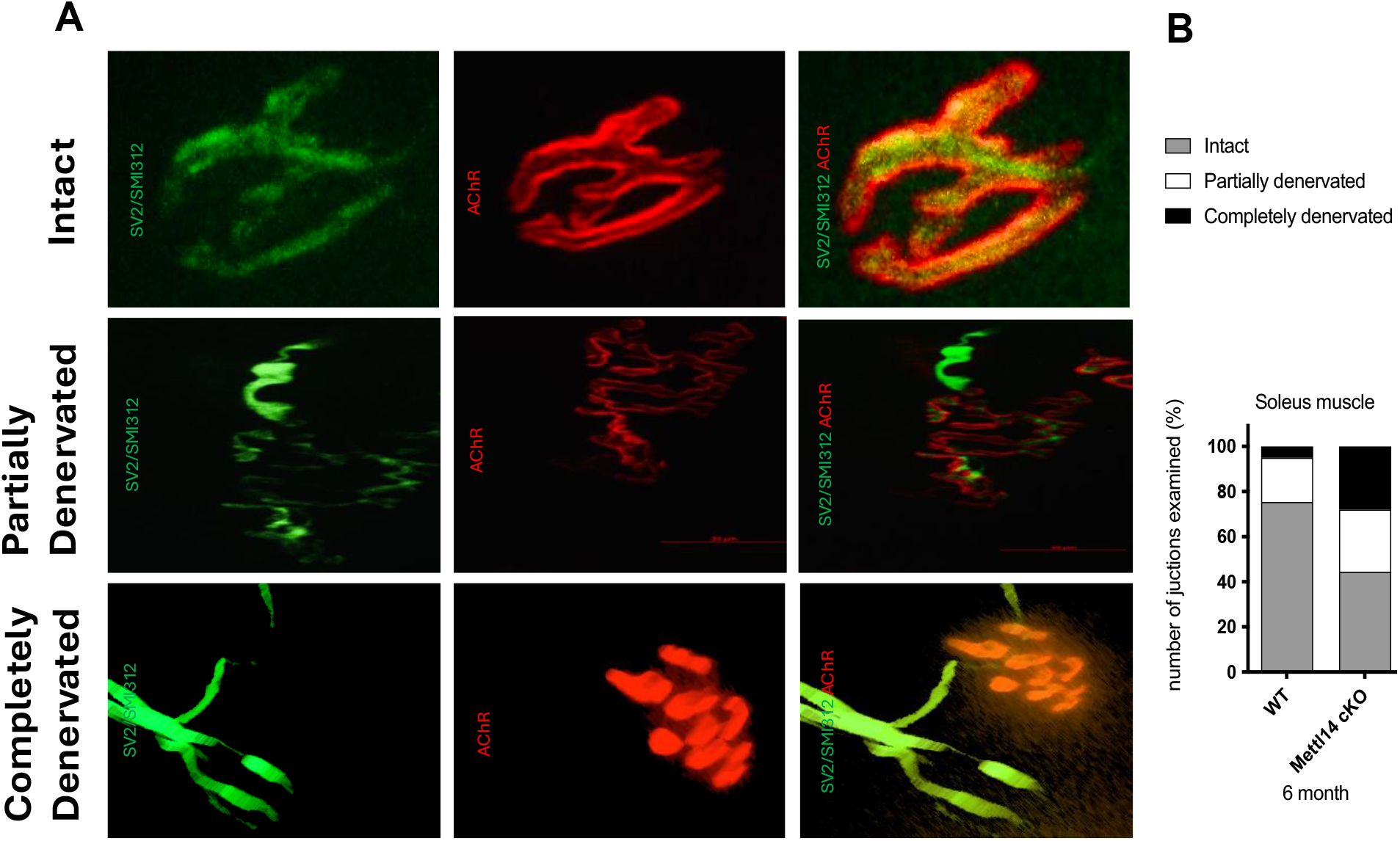
Analysis of neuromuscular junction innervation. Innervation of neuromuscular junctions in the soleus muscle was impaired at 6 month old Mettl14 cKO mice compared with wild type mice. (A) Representative images, scale bar 20 μm. (B) Gray bars indicate intact NMJs, white bars indicate partially denervated NMJs, and the black bars indicate completely denervated NMJs. WT,wild type; Mettl14 cKO, Mettl14 conditional knock out. n= 3 animals per each group.

### Mettl14 cKO mice have impaired peripheral nerve regeneration

To evaluate the impact of Schwann cell-specific Mettl14 deletion on peripheral nerve regeneration, we performed histological analyses of the distal sciatic nerve in wild-type (WT) and Mettl14 cKO mice at 4 months of age—a time point coinciding with the onset of clinical and histological manifestations. A sciatic nerve crush injury was induced in mid-thigh, and tissues were analyzed at 2- and 4-weeks post-injury to assess degree of axonal degeneration and subsequent early regeneration. At 2 weeks post-injury, Mettl14 cKO mice exhibited significantly reduced axonal degeneration compared to WT controls, as quantified by fragmented axons in histological sections (p < 0.01, n=4). This suggests an impaired initial degenerative response. By 4 weeks post-injury, quantification of axonal regeneration profiles revealed that Mettl14 cKO mice exhibited a complete absence of regenerated axons (p < 0.0001, n=3). In contrast, WT mice demonstrated robust regenerative capacity, with significantly higher number of regenerated axon profiles (Figures 4A, B).

**Figure 4.**
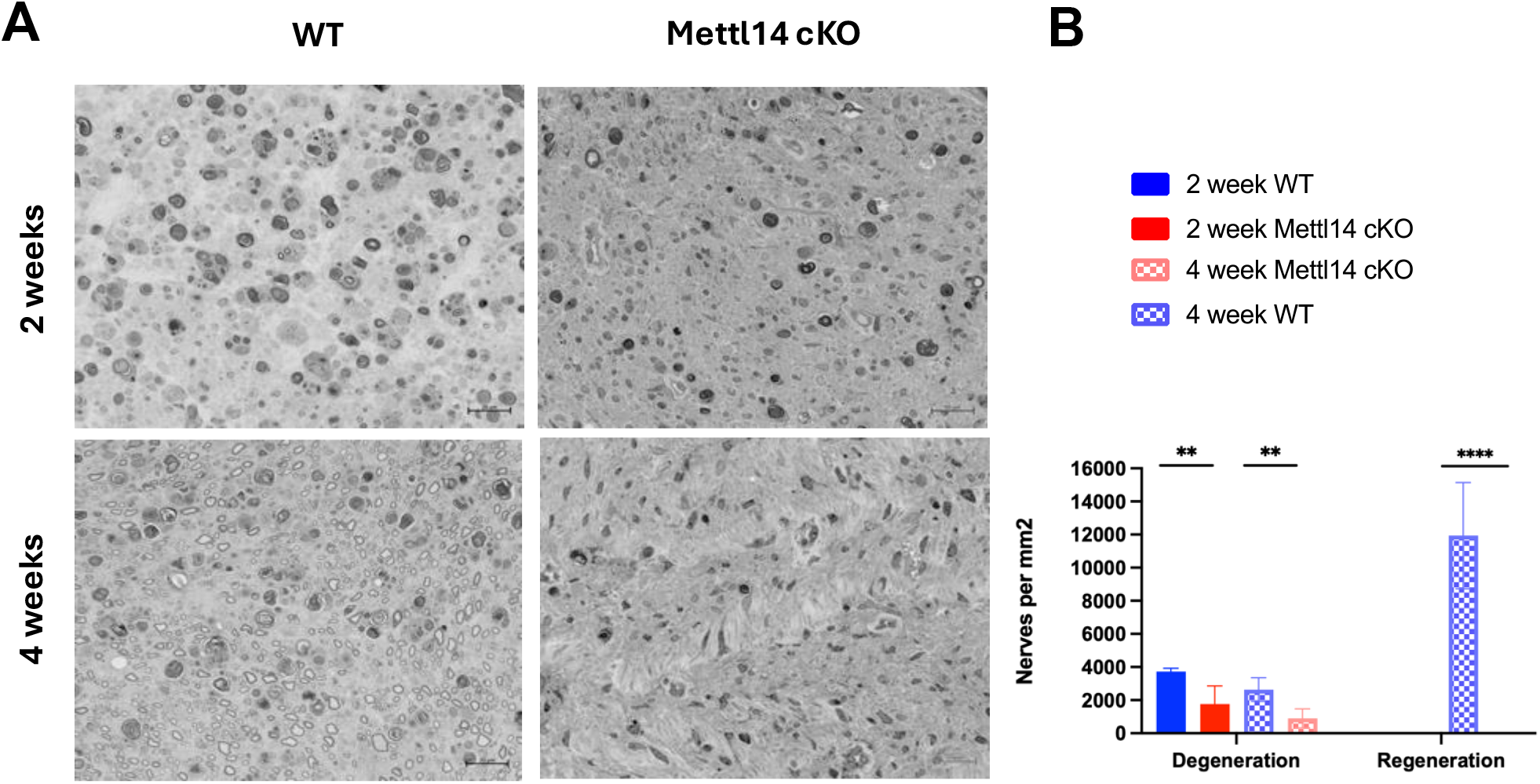
Axonal degeneration and regeneration. Sciatic nerves were harvested from 4-month-old wild-type and Mettl14 cKO mice and plastic sections were prepared for light microscopy. (A) Representative images, scalebar 20μm. Nerve morphometry was performed and (B) degeneration and regeneration profile status are displayed. n=3-4 animals per condition,****p < 0.0001; **p < 0.01. WT,wild type; Mettl14 cKO, Mettl14 conditional knock out.

These findings highlight the essential role of Mettl14 in facilitating efficient axonal regeneration following nerve injury. Specifically, Mettl14 is critical for proper Schwann cell response to denervation and subsequent support of axon regeneration and remyelination and suggested that Mettl14 is important for functional recovery. (Figures 4A, B).

To investigate the mechanisms underlying impaired peripheral nerve regeneration in Mettl14 cKO mice, we examined macrophage recruitment to the site of sciatic nerve injury—a critical process required for myelin debris clearance and subsequent tissue repair (2). We performed these analyses in 4-months of age-a time point coinciding with the onset of clinical and histological manifestations. Immunohistochemical analysis revealed that both wild-type and Mettl14 cKO mice exhibited increased CD68 levels following injury. However, the Mettl14 cKO mice demonstrated a slower rate of macrophage accumulation, attributed to elevated baseline CD68 levels, suggesting impaired macrophage recruitment kinetics in the absence of Mettl14 in Schwann cells. Notably, baseline CD68 positivity was elevated in uninjured sciatic nerves of Mettl14 cKO mice compared to WT controls, indicating ongoing macrophage activation and degeneration even in the absence of acute injury.(Figure 5A,B) Complementary qPCR analysis demonstrated markedly reduced expression of macrophage chemoattractant protein 1 (MCP-1), a key regulator of macrophage recruitment, in Mettl14 cKO sciatic nerves at 1 day post-injury (p < 0.05; Figure 5C).

**Figure 5.**
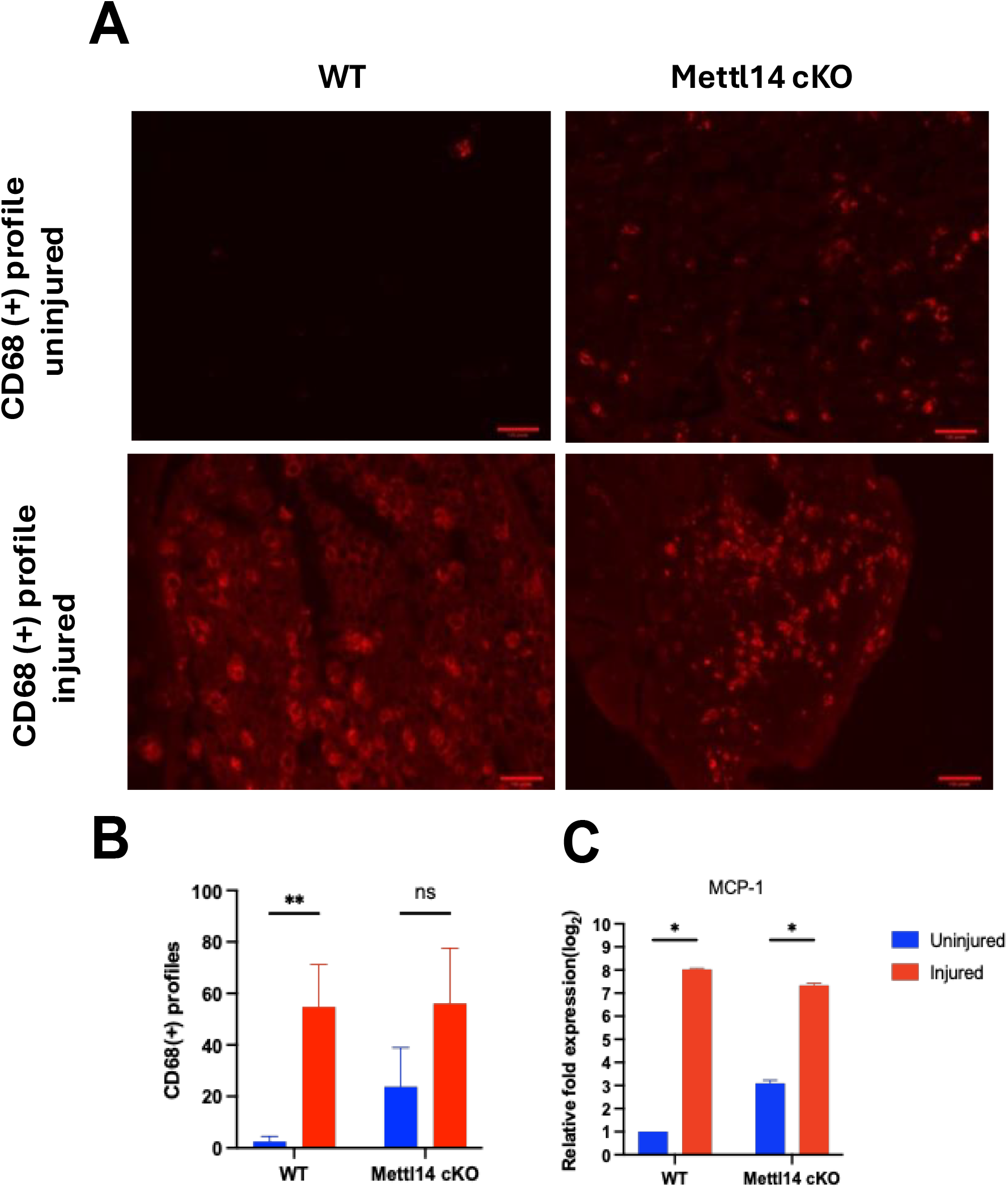
Recruitment of macrophages. Mettl14 may organizes peripheral nerve regeneration by increasing macrophage recruitment to the injured area. Sciatic nerves were harvested from 4-month-old wild-type and Mettl14 cKO mice and processed for immunohistochemistry with an anti-CD68 antibody. (A) Representative images, scale bar 100μm. (B) CD68(+) profile multiple t-tests **p < 0.01; n.s. p > 0.05, (C) qPCR analysis of MCP-1 expression in wild type and Mettl14 conditional knock-out mice following 1 day sciatic nerve transection)****p < 0.0001; unpaired t-test; n=3-4 animals per condition. WT,wild type; Mettl14 cKO, Mettl14 conditional knock out.

### Loss of Mettl14 in Schwann cells impairs cell proliferation and mitochondrial function

To further elucidate the mechanisms underlying regenerative failure in Mettl14 conditional knockout (cKO) mice, we investigated Schwann cell proliferation, a critical process for peripheral nerve repair. (15). Schwann cells derived from neonatal WT and Mettl14 cKO mice were cultured and cell proliferation was assessed using EdU incorporation assays, which measure DNA synthesis during cell division by incorporating the thymidine analog EdU into newly synthesized DNA.(16) These assay revealed a significant decrease in the proportion of proliferating Schwann cells in the Mettl14 cKO group compared to WT controls (p < 0.0001). Notably, Mettl14-deficient Schwann cells exhibited impaired growth and division, leading to proliferative arrest and increased cell death. These findings demonstrate that Mettl14 is indispensable for Schwann cell proliferation and survival, highlighting its broader role in orchestrating the cellular events essential for peripheral nerve regeneration (Figure 6A).

**Figure 6.**
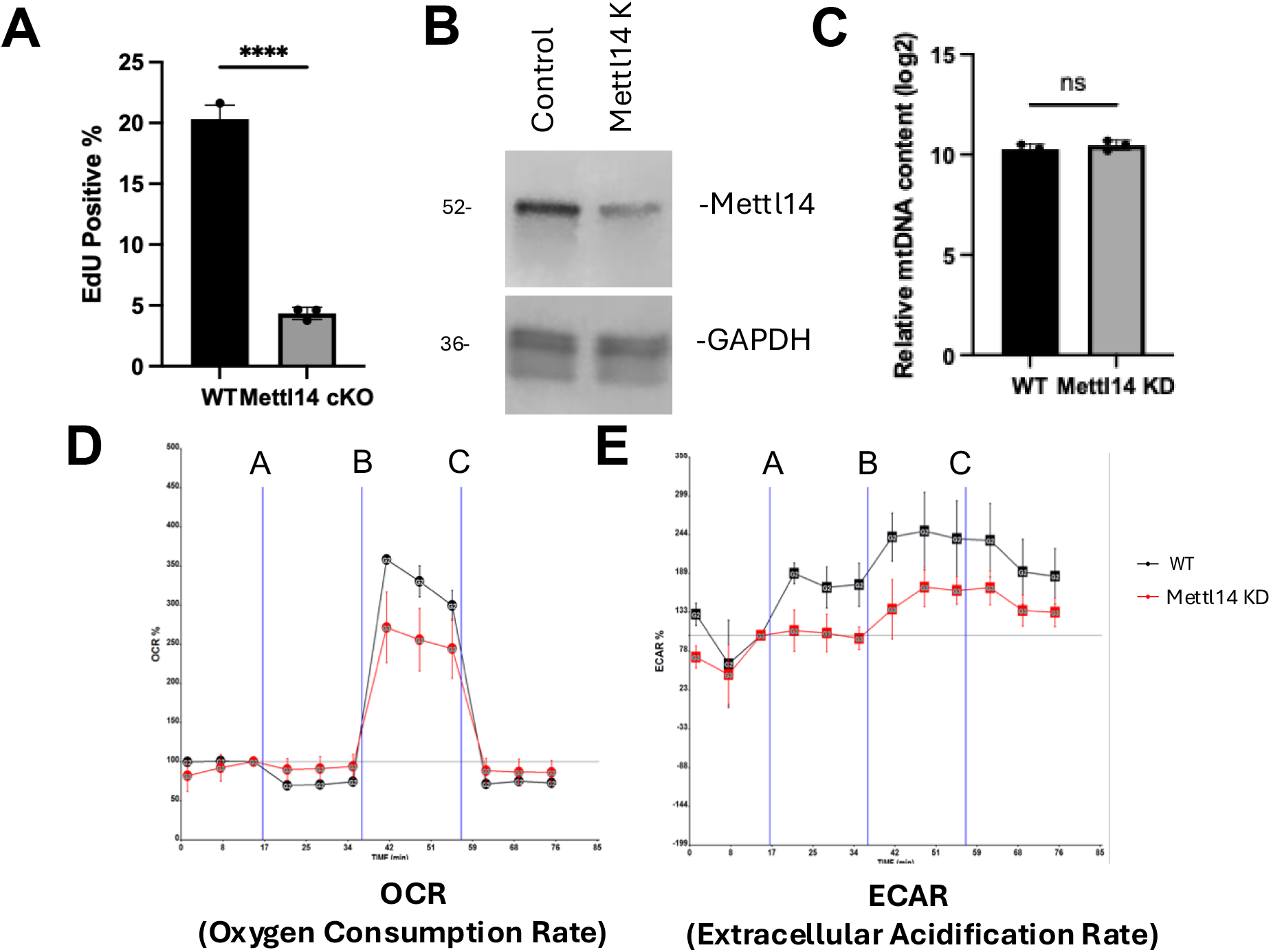
Schwann cell proliferation and mitochondrial function evaluation. Mettl14 cKO Schwann cells exhibit decreased proliferation and impaired mitochondrial function. (A) Percentage of EdU-positive cells, indicating decreased Schwann cell proliferation in Mettl14 cKO cells compared to wild-type. n = 3 animals per condition ****p < 0.0001, unpaired t-test. (B) Western blot analysis showing decreased Mettl14 protein expression in Mettl14 knockdown Schwann cells. (C) Measurement of relative mitochondrial DNA content in Mettl14 cKO Schwann cells. (n.s. p > 0.05) (D) Oxygen consumption rate (OCR) and extracellular acidification rate (ECAR) in Schwann cells seeded in Seahorse assay plates. Mitochondrial bioenergetic function was assessed by sequential injection of (a) 2 μM oligomycin, (b) 4 μM carbonyl cyanide p-(trifluoromethoxy) phenylhydrazone (FCCP), and (c) 0.5 μM rotenone and 4 μM antimycin into the culture media. WT, wild type; Mettl14 cKO, Mettl14 conditional knockout; Mettl14 KD, Mettl14 knock down.

Recognizing the essential role of mitochondria in Schwann cell proliferation and energy metabolism (5), we investigated mitochondrial function in Mettl14 knockdown (KD) Schwann cells. Due to the poor viability of Mettl14 knockout (KO) Schwann cells in primary culture, we knocked down Mettl14 cells by transfecting primary Schwann cells with Mettl14-specific siRNA. Efficient knockdown (KD) of Mettl14 was confirmed by western blot analysis, showing a marked reduction in Mettl14 protein levels compared to controls (Figure 6B). We first quantitatively assessed relative mitochondrial DNA (mtDNA) content using the qPCR method (17), which revealed no significant differences between the WT and Mettl14 KD groups (Figure 6C). Bioenergetic analysis revealed that Mettl14 KD cells exhibited significantly impaired mitochondrial respiration, with reduced maximal oxygen consumption rate (OCR), a measure of mitochondrial oxidative phosphorylation capacity, and spare respiratory capacity, which reflects mitochondrial adaptability under stress (Figure 6D). Furthermore, extracellular acidification rate (ECAR) analysis, an indicator of glycolytic activity, showed normal basal glycolysis but decreased maximal glycolytic capacity and glycolytic reserve (Figure 6E). These findings (Figure 6D,E) suggest that Mettl14-deficient Schwann cells experience bioenergetic impairments, linking mitochondrial dysfunction to defects in proliferation and regeneration. Collectively, this suggests the pivotal role of Mettl14 in regulating Schwann cell bioenergetics and demonstrates how its loss disrupts the cellular mechanisms critical for effective peripheral nerve myelination and repair.

### Transcriptomic analysis

To elucidate the role of Mettl14 in Schwann cell gene expression during myelination and peripheral nerve repair, we performed bulk RNA sequencing (RNA-seq) on sciatic nerves from wild-type (WT) and Mettl14 cKO mice at 4 months of age. Sciatic nerve samples were collected from both intact nerves and injured nerves at three critical time points—1-, 3-, and 7-days post-injury—to capture baseline gene expression as well as the early dynamic transcriptomic changes associated with injury response and repair.

Differential expression analysis revealed distinct gene expression patterns in intact sciatic nerves (Figure 7A), underscoring the pivotal role of Mettl14 in Schwann cell myelination. In intact sciatic nerves from 4-month-old Mettl14 cKO mice, RNA-seq analysis identified significant downregulation of key myelin-associated genes, including Mbp, Pmp22, and Mpz, which are essential for proper myelin formation and maintenance (18). This downregulation is consistent with the beginning of demyelination observed in these mice at this time point.

**Figure 7.**
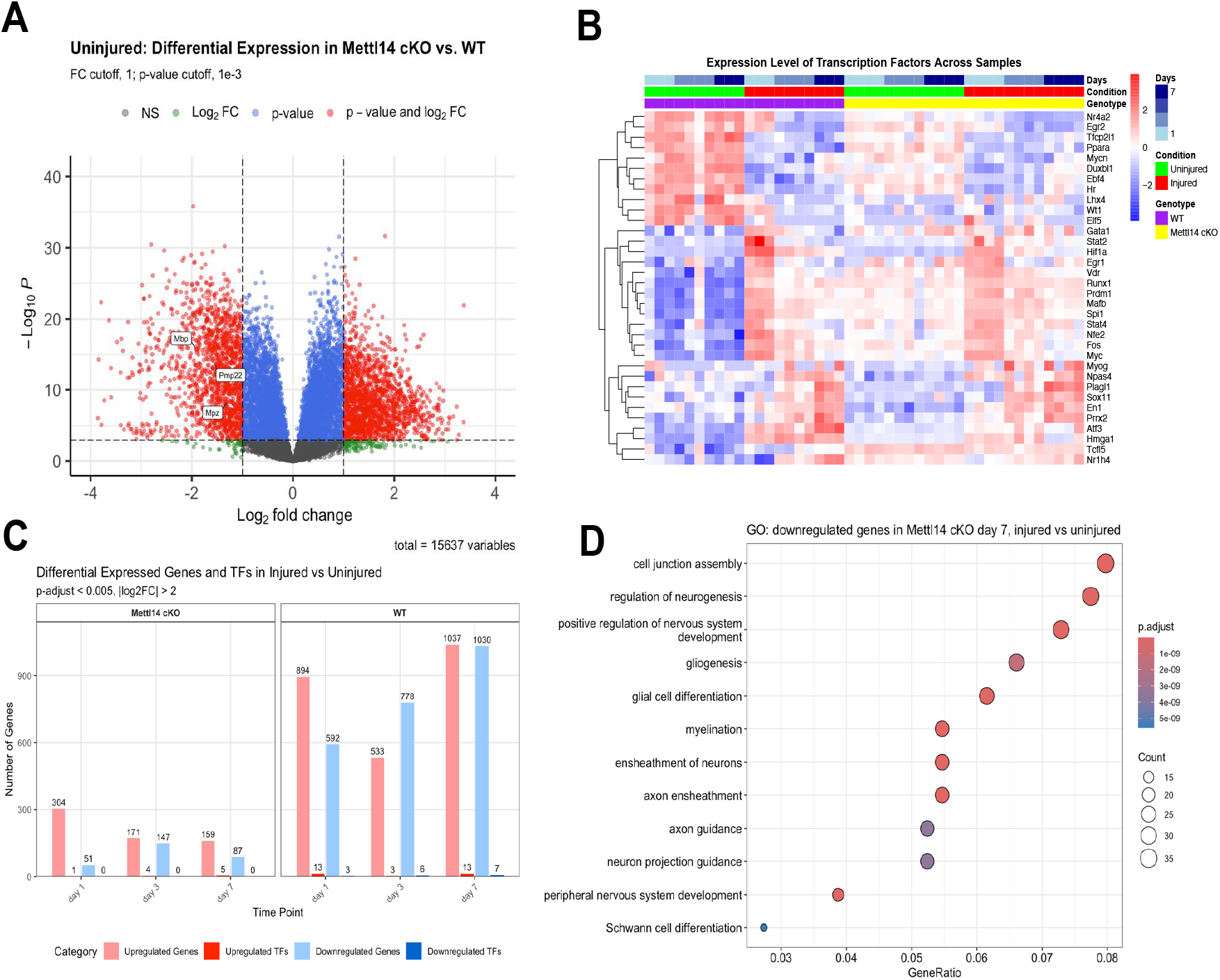
Transcriptomics analysis Figure 7A. Volcano plot depicting differentially expressed genes in Mettl14 cKO mice compared to WT under uninjured conditions. Each point represents a gene, with color indicating statistical significance and fold change criteria: non-significant (NS, gray), significant by fold change alone (|log2FC| > 1, green), significant by p-value alone (p < 0.001, blue), and significant by both fold change and p-value (|log2FC| > 1 and p < 0.001, red). Select myelin-related genes (Mbp, Pmp22, Mpz) are labeled. **Figure 7B. Heatmap illustrating the scaled expression levels (row-normalized, variance-stabilized counts) of transcription factors across samples from Mettl14 cKO and WT mice**. Rows represent individual transcription factors, while columns correspond to samples, grouped by genotype (Mettl14 cKO vs. WT), condition (Uninjured vs. Injured), and time points (Days 1, 3, and 7). Expression values are mean-centered and scaled per gene (row), where red indicates expression above the gene’s mean across all samples, and blue represents below-mean expression. Key transcription factors are labeled on the right. **Figure 7C. Differential gene expression analysis comparing injured versus uninjured conditions in Mettl14 cKO and WT mice**. Bar plot showing the number of differentially expressed genes and transcription factors (TFs) at days 1, 3, and 7 post-injury. Light red and dark red bars represent upregulated genes and TFs, respectively, while light blue and dark blue bars represent downregulated genes and TFs, respectively. Differential expression was defined by adjusted p-value < 0.005 and |log2 FC| > 2. Numbers above bars indicate the exact count of differentially expressed genes or TFs in each category. The data are separated into two panels showing the results for Mettl14 conditional knockout (cKO) and wild-type (WT) mice. **Figure 7D. Gene Ontology (GO) analysis of downregulated genes in Mettl14 conditional knockout mice at day 7 post-injury compared to uninjured group**. Dot plot showing significantly enriched GO terms for biological processes among downregulated genes (log2FC < -1, adjusted p < 0.05) in injured versus uninjured tissue from Mettl14 conditional knockout mice. The analysis reveals enrichment of processes related to neuronal development, myelination, and glial cell function. GO terms are arranged by decreasing GeneRatio values.

Following nerve injury, Mettl14 cKO mice exhibited severe transcriptomic dysregulation, particularly at 7 days post-injury. Genes involved in axonogenesis and repair, such as Egr2 and Atf3, were markedly downregulated in Mettl14 cKO nerves compared to WT controls. A heatmap of transcription factor expression further revealed that injury-responsive genes, including Stat2 and Atf3, were robustly upregulated in WT mice but blunted or absent in Mettl14 cKO mice (Figure 7B, C). Functional pathway analysis revealed a striking downregulation of key signaling cascades involved in myelin assembly and axon guidance (Figure 7D), indicating that Mettl14 is indispensable for orchestrating the genetic programs that drive nerve regeneration.

Together, these transcriptomic insights establish Mettl14 as a central regulator of Schwann cell gene expression, governing both myelination and injury-induced transcriptional reprogramming. Loss of Mettl14 disrupts essential transcriptional networks, resulting in impaired nerve repair, defective myelination, and progressive phenotypic deficits in Mettl14 cKO mice. These findings position Mettl14 as a key molecular driver of Schwann cell maintenance and peripheral nerve regeneration, highlighting its potential as a therapeutic target for demyelinating neuropathies.

## Discussion

RNA modifications, especially N6-methyladenosine (m6A), have been identified as key post-transcriptional regulatory mechanisms that significantly influence gene expression (19). Among the various types of RNA modifications, m6A is the most common internal modification found in eukaryotic mRNA (20). Notably, m6A methylation differs from DNA and protein methylation in its ability to rapidly induce transcriptome changes, especially during critical cell-state transitions (19). These rapid modifications of the m6A landscape allow for swift and flexible changes in cellular phenotypic properties, enabling cells to adapt quickly to environmental and developmental cues (12,21). In the peripheral nervous system (PNS), such dynamic regulation is crucial, particularly during processes like myelination, nerve injury response, and regeneration, where the ability of Schwann cells to rapidly adjust gene expression is essential for maintaining nerve function (22).

Our study suggests that N6-methyladenosine (m6A) RNA methylation, regulated by Mettl14, does not appear to play a role in Schwann cell development but have a crucial role in maintaining Schwann cell function, myelination, and peripheral nerve regeneration. Specifically, Mettl14 deletion in Schwann cells leads to progressive demyelination characterized by demyelinating peripheral neuropathy and impaired axonal regeneration. While developmental myelination proceeds normally, myelin integrity deteriorates over time, with Mettl14 cKO mice displaying reduced myelin thickness and axon density by four months and severe demyelination by 12 months. These findings underscore the indispensable role of m6A methylation in sustaining peripheral nerve integrity and function.

This role differs from established findings in the central nervous system (CNS), where m6A methylation regulates oligodendrocyte maturation and myelination. For instance, Mettl14 is essential for oligodendrocyte development, and its loss disrupts transcriptome regulation and leads to CNS hypomyelination (13). Similarly, Prrc2a, an m6A reader, stabilizes Olig2 mRNA to ensure oligodendrocyte specification and myelination, with its absence resulting in severe hypomyelination and neurological defects (23). Extending these findings to the peripheral nervous system (PNS), our study highlights a unique aspect: while Mettl14-mediated m6A methylation is nonessential for Schwann cell development, it is critical for long-term stability, myelin maintenance, and regenerative function. The age-dependent degeneration of large, myelinated axons, critical for proprioception and motor function, likely explains the progressive motor deficits observed in Mettl14 cKO mice. (24) Hindlimb motor impairments, detected by three months, precede forelimb deficits, consistent with the increased vulnerability of longer sciatic nerves. (25,26).

Our findings, combined with prior studies, underscore the critical role of m6A methylation as a dynamic regulator of axonal regeneration and neuronal survival. Notably, the m6A demethylase ALKBH5 has been identified as an inhibitor of axonal regeneration, with its knockdown shown to enhance regenerative capacity by modulating lipid metabolism-related genes such as Lpin2 (27). Dysregulation of m6A pathways, whether through writers like METTL3 and METTL14 or erasers like FTO and ALKBH5, has been implicated in neurodegenerative processes and impaired regenerative responses (28).

Macrophages are recruited to the injury site to phagocytose myelin and axonal debris, a process critical for creating a conducive environment for nerve regeneration. This recruitment occurs before and during Schwann cell proliferation (29,30). Reduced macrophage recruitment leads to impaired sensory axon regeneration (31). Our findings of reduced macrophage recruitment further emphasize the critical role of m6A methylation in orchestrating immune responses essential for effective nerve regeneration. The impaired macrophage response observed in Mettl14 cKO mice, combined with their increased baseline macrophage activity, highlights the dual role of Mettl14 in maintaining nerve homeostasis and orchestrating the cellular and molecular events that drive regenerative inflammation (32). Together, these results suggest a novel link between Mettl14-mediated regulation of inflammation and the functional recovery of injured peripheral nerves.

A significant finding of our study is the critical impact of Mettl14 deletion on Schwann cell survival and proliferation. The inability to sustain the growth of Mettl14-deficient Schwann cells in culture highlights the essential role of m6A methylation in maintaining Schwann cell viability and proliferative capacity. This defect likely contributes to the regenerative failure observed in Mettl14 cKO mice, as Schwann cell proliferation is fundamental to peripheral nerve repair following injury. The underlying mechanisms of this proliferation defect warrant further investigation.

Mitochondria are crucial for bioenergetic metabolism and mitochondrial dysfunction can lead to oxidative stress, energy deficiency, and imbalances in mitochondrial dynamics, all of which contribute to axon degeneration and regeneration (5,27). In diabetic neuropathy, for instance, mitochondrial dysfunction is linked to a shift from oxidative phosphorylation to anaerobic glycolysis, impairing nerve growth and regeneration (33). Mitochondrial dysfunction in SCs can lead to nerve conduction abnormalities, demyelination, and axonal degeneration. Ensuring the proper function of SC mitochondria is essential for maintaining peripheral nerve health and facilitating regeneration (5,27). Our results suggest that m6A methylation is essential for maintaining Schwann cell metabolism, under conditions of high metabolic demand during nerve injury and repair.

Despite the significant findings of our study, several limitations should be acknowledged. First, while we demonstrated that m6A methylation is essential for Schwann cell function and nerve regeneration, the precise molecular mechanisms underlying these effects remain incompletely understood. Second, our study relied exclusively on animal models, which, while invaluable for mechanistic insights, may not fully recapitulate the complexity of human Schwann cell biology and peripheral nerve regeneration. Bridging this gap will require additional studies using human-derived cells or tissues. Third, although we identified defects in Schwann cell proliferation in vitro, it remains unclear whether these are primarily cell-autonomous effects or arise from altered interactions with other cell types or the extracellular microenvironment. Additionally, our focus on the motor and sensory components of the peripheral nervous system did not extend to a detailed analysis of the autonomic or enteric nervous systems. Notably, the increased incidence of rectal prolapse in aged Mettl14 cKO mice suggests potential involvement of Schwann cells in autonomic or enteric nervous system dysfunctions. Future studies are needed to explore these intriguing possibilities and the broader implications of Mettl14 deletion in non-myelinating Schwann cells. By addressing these limitations in future research, we can deepen our understanding of m6A methylation in Schwann cell biology and uncover its full therapeutic potential in peripheral nerve disorders.

In summary, our findings suggest that targeting the m6A machinery, including Mettl14, offers a promising therapeutic avenue. By modulating m6A pathways, it may be possible to enhance Schwann cell-mediated myelination, repair and address the multifaceted challenges of nerve regeneration and neurodegeneration.

## Materials and Methods

### 1.1 Animals and Housing

All experiments were approved by the Johns Hopkins Animal Care and Use Committee. Mettl1 ^flox/flox^ mice were provided by Dr. Guo Li Ming (Perelman School of Medicine, University of Pennsylvania), and P0-Cre mice were obtained from Jackson Laboratory [catalogue # 017927]. Mettl14^flox/flox^ mice were crossed with P0-Cre mice to Mettl14 conditional knockout (cKO) mice for experimentation. P0-Cre x Mettl14^flox/WT^ heterozygous littermates were used as controls. Mice were housed in a single room maintained at 72 ± 5°F and 42% relative humidity, with a 12-hour light/dark cycle (lights on at 06:30 h). Animals were housed in same-sex littermate groups and fed a standard mouse diet with ad libitum access to water.

### 1.2 Genotyping

Mouse genomic DNA was extracted from ear tissue and analyzed via PCR to detect Cre and Mettl14 floxed alleles (Supplemental Table 1). PCR amplification was performed using a master mix (Invitrogen, CA, USA) according to the manufacturer’s instructions. For Cre genotyping, the conditions were: 94 °C for 3 min, at 94 °C for 30 sec, at 51.7 °C for 1 min followed by 35 cycles at 72 °C for 1 min, at 72 °C for 5 min. For Mettl14 genotyping, the conditions were: 94 °C for 3 min, 94 °C for 30 sec, 56 °C for 30 sec followed by 35 cycles of 72 °C for 30 s, 72 °C for 5 min. PCR products (100 bp Cre transgene and 319 bp Mettl14 floxed allele) were seperated by 1.5% agarose gel electrophoresis and visualized by Bio-Rad Universal Hood II Gel Imaging System.

## 2. Functional Tests

Mice were assessed monthly from 3-4 weeks to 12 months by a single blinded investigator to minimize inter-examiner variability. The investigator was unaware of the animal genotypes during assessments. Functional tests included the neuromuscular SHIRPA protocol, accelerated Rotarod, and grip strength assays (34). Nine age-separated groups of mice were used. After each test, equipment was cleaned with 10% ethanol and dried with paper towels.

### 2.1 Neuromuscular SHIRPA

The neuromuscular function of the mice was assessed using a modified SHIRPA protocol (35). Fourteen non-invasive tests were performed to evaluate body posture, spontaneous movement, and tremor in a viewing jar. Transfer arousal, gait, pelvic elevation, tail elevation, and touch escape responses were assessed in an open arena. Additional tests, including trunk curl, limb grasping, toe pinch, and hanging grasp, were performed while the mouse was suspended above the arena. The SHIRPA screen provided a detailed assessment of neuromuscular and physical function, with scores ranging from 0 (no impairment) to 49 (severe impairment). Scores are positively correlated with clinical impairment.

### 2.2 Accelerated Rotarod Test

Ataxia, balance, and motor coordination were assessed using an accelerated Rotarod (36) over three consecutive days. Mice were placed on a rotating drum that began at 4 rpm and increased by 0.5 rpm every 30 seconds, with a maximum speed of 40 rpm. Each mouse underwent nine trials after an initial training session. Data were recorded for latency to fall (in seconds) and maximum velocity (in rpm). Standard fluorescent lighting was used in the testing room, and mice were returned to their home cages between trials.

### 2.3 Stimulated Grip Strength Tests

Forelimb and hindlimb grip strength were measured using a force-transducer apparatus (37). Testing was performed in three trials separated by five-minute rest intervals to prevent fatigue. Mice were held by the tail and lowered onto a metal grid, which they were allowed to grasp. The mice were gently pulled backward to measure the peak pull force (in grams). The results from all trials were averaged for data analysis.

## 3. Nerve Morphometry

Sciatic nerves were collected bilaterally from both wild-type and Mettl14 cKO mice, immediately fixed, and processed for electron microscopy following established methods (4). Tiff images of sciatic nerve sections were captured using a Zeiss Apotome 3 microscope equipped with a 63X oil objective. Semi-thin sections stained with toluidine blue were used to count the total number of myelinated axons per cross-section (38). Axons were analyzed from at least four randomly selected rectangular areas of equal size within each sciatic nerve. Myelin sheaths and axons were manually traced and counted using ImageJ with the appropriate plugin. (http://gratio.efil.de). Axon diameters and g-ratios were calculated based on the areas of the axon and the combined axon-plus-myelin sheath. Total axon counts were estimated by determining the total area of the sciatic nerve and applying a selected area-to-total area ratio to extrapolate the axon counts. At least 200 axons per mouse were evaluated.

## 4. Analysis of neuromuscular junctions

Soleus muscles were post-fixed in 4% paraformaldehyde, washed in PBS with 0.1 M glycine, and incubated with rhodamine-conjugated α-bungarotoxin (Invitrogen) to stain acetylcholine receptors. Tissues were then permeabilized in cold methanol and blocked with 0.2% Triton and 2% BSA. Overnight incubation with primary antibodies, SMI 312 (Sternberger Monoclonals) and SV2 (Developmental Studies Hybridoma Bank), was performed to label axons and nerve terminals, respectively. After washing, secondary antibodies were applied, and the tissues were mounted in Vectashield [catalogue # H-1200] (Vector Laboratories). Neuromuscular junctions (NMJs) were categorized as intact, fully denervated, or partially denervated based on the overlap between nerve terminals and postsynaptic acetylcholine receptors. At least 50 NMJs per muscle were analyzed, with three animals per group.

To evaluate denervation, each neuromuscular junction was first categorized as follows: (1) completely intact as determined by 100% overlap between the nerve terminal arbor and postsynaptic AChRs; (2) completely denervated, as identified by the lack of any nerve terminal labeling at identified postsynaptic AChRs or (3) partially denervated as determined by the presence of less than 100% overlap between nerve terminals and postsynaptic AChRs. A minimum of 50 neuromuscular junctions (NMJs) were analyzed per muscle, with three animals per group.

## 5. Axonal Degeneration and Regeneration Analysis in Sciatic Nerve Histology

To assess axonal degeneration and regeneration, a unilateral sciatic nerve crush injury was performed in mid-thigh section and animals were allowed to recover. Tow and 4 weeks later, distal sciatic nerves were collected from both wild-type and Mettl14 cKO mice. Contralateral sides were collected as controls. The samples were immediately fixed and processed for electron microscopy following established protocols (4). Quantitative analysis was conducted by counting degenerating and regenerating axons in five randomly selected fields per section, with averages calculated across sections from a minimum of three animals per group to ensure robust statistical validity (38).

## 6. CD-68 Immunofluorescent Staining

Distal sciatic nerve immunohistochemistry samples were prepared and processed as described (39). Briefly, the nerves were dissected and incubated overnight in 4% PFA, and dehydrated in 30% sucrose in 1× PBS prior to embedding in Tissue-Tek® O.C.T. compounds (Sakura Finetek, #4583, Torrance, CA, USA). The samples were cryo-sectioned at 15 μm. For diminishing non-specific binding, the samples were first blocked in 5% normal goat serum[catalogue#005-000-021] Jackson Laboratories in 1× PBS for 1 h at RT after permeabilizing with 0.2% TrixonX-100 for 10 min. As primary Antibody CD68 [catalogue#14-0681-82 (Goat anti-Rat, 1:250) (Invitrogen, PA5-109344, Waltham, MA, USA) used. Samples were incubated overnight at 4 °C with it. Respective secondary antibody (405 nm, anti-rat IgG (H + L) (1:1000) (Invitrogen, A48261, Waltham, MA, USA) for 1 h at room temperature. Mounted (ProLongTM Gold Antifade Mountant, Invitrogen, P36930), imaged under Zeiss Apotome 3 and analyzed under Image J.

## 7. Schwann cell culture and EdU cell proliferation assay

Schwann cells were isolated from the sciatic nerves of 3-day-old neonatal pups according to established protocols as described previously (40). The harvested cells were cultured in DMEM [catalogue # 11965-0902] (Invitrogen, Carlsbad, CA) containing 10% FBS [catalogue# 26140079](Invitrogen), 1% penicillin-streptomycin (Invitrogen), and 2 μM forskolin [catalogue # f6886], (Sigma, St. Louis, MO), in a humidified incubator with 5% CO2 at 37°C. Schwann cell proliferation was measured using the Click-iT® EdU Imaging Kit (ThermoFisher Scientific). Briefly, cells were seeded in poly-L-lysine-coated 96-well plates, and EdU (20 μM, ThermoFisher Scientific) was added to the culture for 24 hours. Afterward, cells were fixed with 4% paraformaldehyde in phosphate-buffered saline (PBS) and stained with Hoechst 33342. Fluorescent images were acquired using a microscope, and Schwann cell proliferation was calculated by dividing the number of EdU-positive cells by the total cell count.

## 8. Schwann cell culture and Seahorse bioenergetic analysis

Schwann cells were cultured from the sciatic nerves of 4-week-old mice, with minor modifications from a previously described protocol (4). Briefly, the sciatic nerves were dissected, cut into small segments, and enzymatically dissociated using 0.3% collagenase[catalogue#17100-017] (Gibco) for 45 minutes, followed by 0.25% trypsin for 15 minutes. Dissociated cells were resuspended in Schwann cell medium (ScienceCell), which contains basal media supplemented with 10% FBS, 1% Schwann cell growth supplement, and 1% penicillin/ streptomycin[catalogue # 15070063]. On the first day post-plating, the medium was replaced with Schwann cell purification media, containing 10 μM arabinosylcytosine [catalogue#147-94-4] (AraC, Millipore Sigma) for two days. The media was alternated between Schwann cell media and purification media for four more days. Bioenergetic analysis was performed using the XF96 extracellular flux analyzer (41) by sequentially injecting 2 μM oligomycin, 4 μM FCCP, 0.5 μM rotenone, and 4 μM antimycin. The oxygen consumption rate (OCR) and extracellular acidification rate (ECAR) were recorded.

## 9. RNA Isolation and qPCR Experiment

Total RNA was extracted from sciatic nerves using Trizol reagent [catalogue # 15596026, Invitrogen], following the manufacturer’s protocol. cDNA synthesis was performed using the QuantiTect Reverse Transcription Kit (Qiagen). Quantitative real-time PCR (qPCR) was conducted using Sybr-Green[catalogue # KCQS00] to measure RNA levels. Relative gene expression was normalized to Peptidylprolyl Isomerase A (PPIA). The primer sequences are listed in Supplemental Table 1. Gel electrophoresis was used to confirm the correct product size and the absence of nonspecific bands. The fold change in expression was calculated using the delta-delta Ct method.

## 10. Protein Isolation and Western Blotting

Sciatic nerves were harvested, homogenized in ice-cold lysis buffer (RIPA Buffer, catalogue#R0278 Thermo Scientific), and centrifuged at 10,000 × g for 15 minutes. The supernatant was stored at -80°C. Protein concentrations were measured using the Pierce BCA Protein Assay Kit[catalogue#23228]. Equal amounts of protein were loaded into SDS polyacrylamide gels (4–15%, BioRad), transferred onto nitrocellulose membranes, and probed with anti-MBP antibodies. Bands were visualized with enhanced chemiluminescence, and membranes were stripped and reprobed for Beta Tubulin and GAPDH, as appropriate.

## 11. RNA-Seq Data Processing and Analysis

Bulk RNA sequencing (RNA-seq) was performed on sciatic nerve samples, with libraries prepared and sequenced at the UCLA Genomics Facility using an Illumina sequencing platform. Raw sequencing reads underwent quality control (QC) assessment using FastQC, and adapters were trimmed using Trim Galore. High-quality reads were then aligned to the mouse reference genome (GRCm39/mm39) using STAR (Spliced Transcripts Alignment to a Reference) with default parameters optimized for mammalian transcriptomic data. To quantify transcript abundance, we used Salmon, which estimates transcripts per million (TPM) values based on quasi-mapping and fragment length distribution. To ensure robust and biologically meaningful downstream analysis, several stringent filtering steps were applied. Expression data were log2-transformed to improve visualization and normalize variance across samples. We removed outliers based on principal component analysis (PCA) and hierarchical clustering to eliminate samples with abnormal transcriptomic profiles. To exclude lowly expressed genes that could introduce noise, we applied a TPM > 0.5 threshold in at least 80% of the samples, retaining only genes with sufficient expression levels for differential expression analysis. Differential gene expression analysis was performed using linear-regression model accounting for batch effects, with q-values calculated as false discovery rate (FDR)-adjusted p-values using the FDR correction. Expression values, fold changes, and statistical significance were adapted for visualization using a log2 transformation, allowing for clearer interpretation of transcriptional dynamics across conditions. (42,43,44,45,46,47,48,49)

## 12. Statistical analysis

All experiments involved two groups: wild-type and Mettl14 cKO. Statistical analyses were performed using GraphPad Prism 10 software. Significance was determined using unpaired Student’s t-tests or two-way ANOVA with Sidak’s multiple comparisons test, as appropriate. A p-value of <0.05 was considered statistically significant.

## Supporting information

Metadata

4 months Mettl14 cKO movie 1

4 months Mettl14 cKO movie 2

4 months Mettl14 cKO movie 3

12 months Mettl14 cKO movie 1

12 months Mettl14 cKO movie 2

12 months Mettl14 cKO movie 3

## Acknowledgments

We would like to acknowledge the Johns Hopkins University School of Medicine Behavioral Core for providing guidance and training for behavioral assays.

## Declaration of generative AI and AI-assisted technologies in the writing process

I declare that first author have utilized OpenAI’s ChatGPT-4, an artificial intelligence language model, to assist in the development of this work. The model was used for language editing. After using this tool, the authors reviewed and edited the content as needed and take full responsibility for the content of the publication.

## Author contributions

Conceptualization: AH

Methodology: AH

Investigation: MCS, AJ, ATY, XH, RK, TH

Visualization: MCS, RM, WC

Supervision: AH

Writing—original draft: MCS, AH

Writing—review & editing: MCS, AJ, ATY, XH, TH, WC, RK, ST, VS, GLM, AH

## Data and materials availability

All data are available in the main text or the supplementary materials.

