## Supplementary Materials for "Lack of mRNA Methylation in Schwann Cells Results in Demyelination and Regenerative Failure"

Mehmet Can Sari *et al.*

**This PDF file includes:**

Figs. S1 to S3
Tables S1
Movies S1

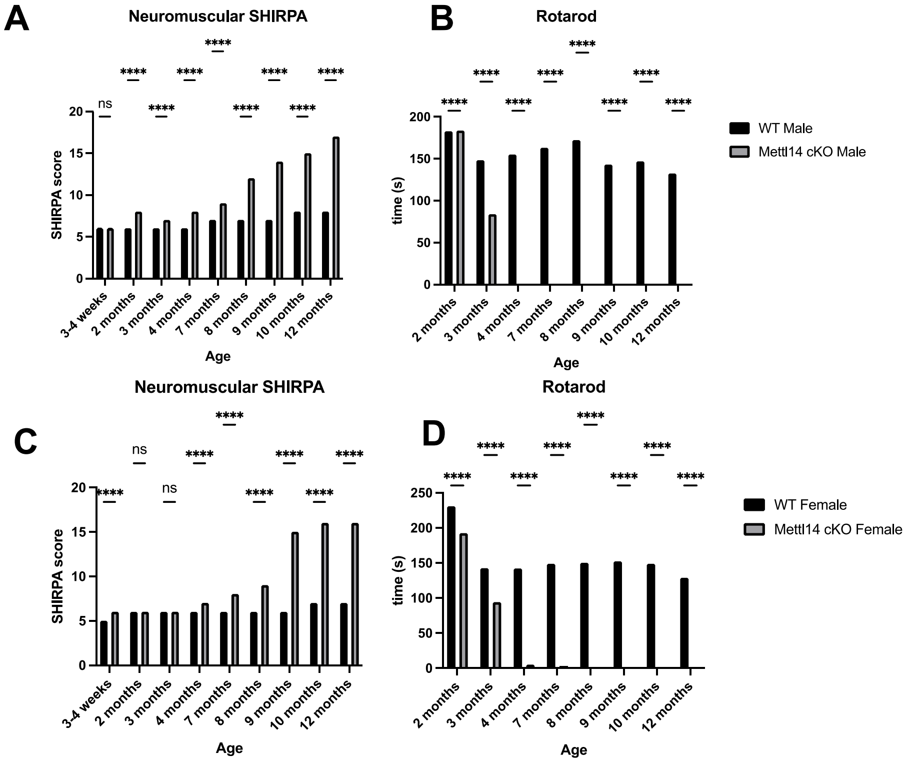


**Figure S1. Clinical characterization of male and female mice analyzed separately on Neuromuscular SHIRPA and Rotarod performance**

3-4 weeks to 12 month- old wild type and Mettl14 cKO mice were evaluated by (A) Neuromuscular SHIRPA, (B) accelerating rotarod assay, n= 3-20 animals per each time point group, two-way ANOVA, Sidak test, ****$p < 0.0001$ ; ns (non significant) $p > 0.05$. WT,wild type; Mettl14 cKO, Mettl14 conditional knock out.

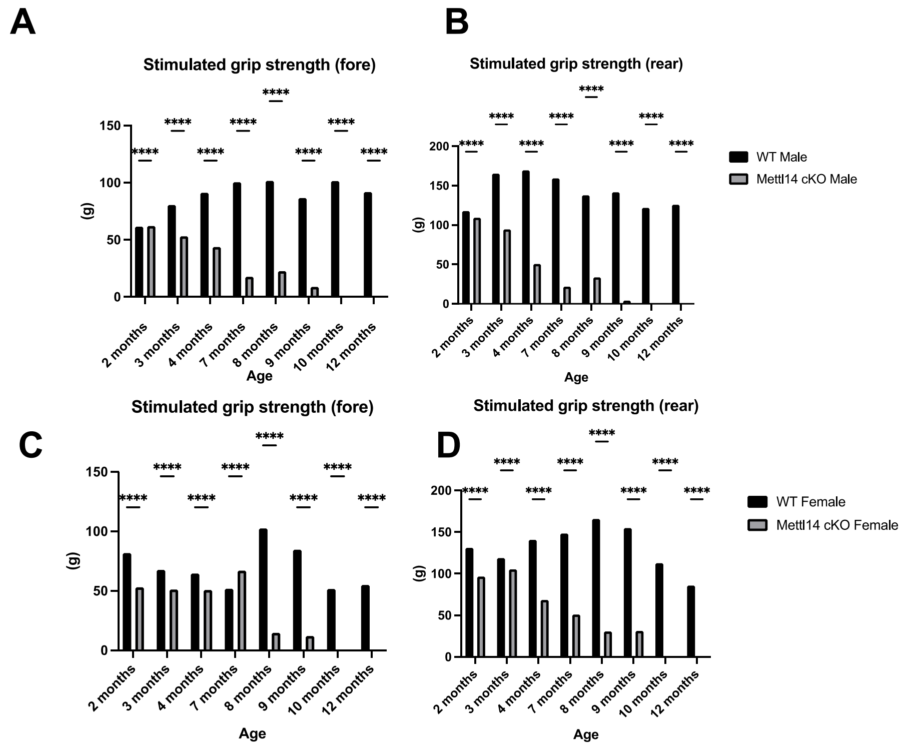


**Fig. S2. Clinical characterization of male and female mice analyzed separately on forelimb grip strength and hidlimb grip strength tests**

3-4 weeks to 12 month- old wild type and Mettl14 cKO mice were evaluated by (A) Neuromuscular SHIRPA, (B) accelerating rotarod assay, n= 3-20 animals per each time point group, two-way ANOVA, Sidak test, ****p < 0.0001. WT,wild type; Mettl14 cKO, Mettl14 conditional knock out.

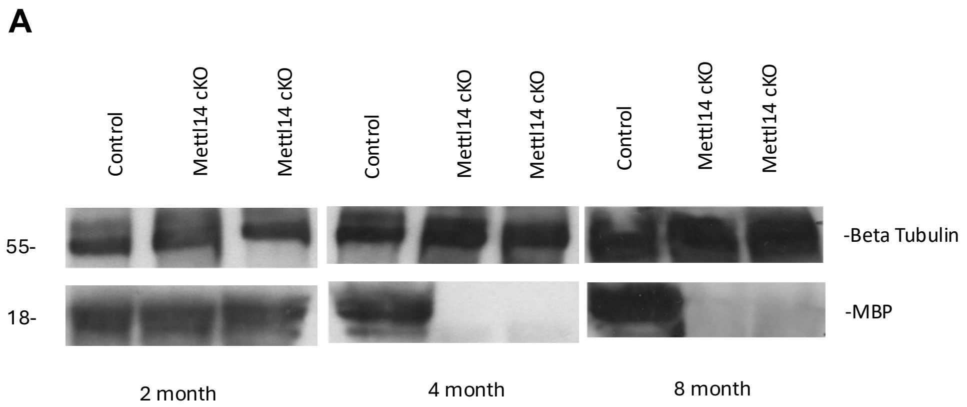


**Fig. S3. Western blot analysis of Beta Tubulin and MBP expression in control and Mettl14 cKO mice at different ages**

(A) Representative Western blot images showing the expression of Beta Tubulin (55 kDa) and Myelin Basic Protein (MBP; ~18 kDa) in the control and Mettl14 conditional knockout (cKO) mice sciatic nerves at 2 months, 4 months, and 8 months of age. Beta Tubulin used as a loading control.

| Primers | Sequence |
|---|---|
| Cre Forward | CGC GTC TGG CAG TAA AAA CTA TC |
| Cre Reverse | GTG AAA CAG CAT TGC TGT CAC TT |
| PosCre Forward | CTA GGC CAC AGA ATT GAA AGA TCT |
| PosC Reverse | GTA GGT GGA AAT TCT AGC ATC ATC C |
| Mettl14 Common | ACT TCA CTT CCA ACG CAG GT |
| Mettl14 Deleted | AGG TTT GTG AAT TCT GAC GCA |
| Metll14 Floxed | ACA TGC TAG AGT CAG CAC CT |
| MCP-1 Forward | AGG TCC CTG TCA TGC TTC TG |
| MCP-1 Reverse | TGG GAT CAT CTT GCT GGT GA |

**Table S1. List of primers and their sequences used for genotyping and gene expression analysis**

Primers listed include sequences for detecting Cre recombinase (Cre Forward and Reverse), positive Cre control (PosCre Forward and Reverse), and Mettl14-specific alleles (Common, Deleted, and Floxed). Additionally, primers for MCP-1 (monocyte chemoattractant protein-1) expression analysis are included.

Age — Mettl14 cKO mice videos

4 months

12 months

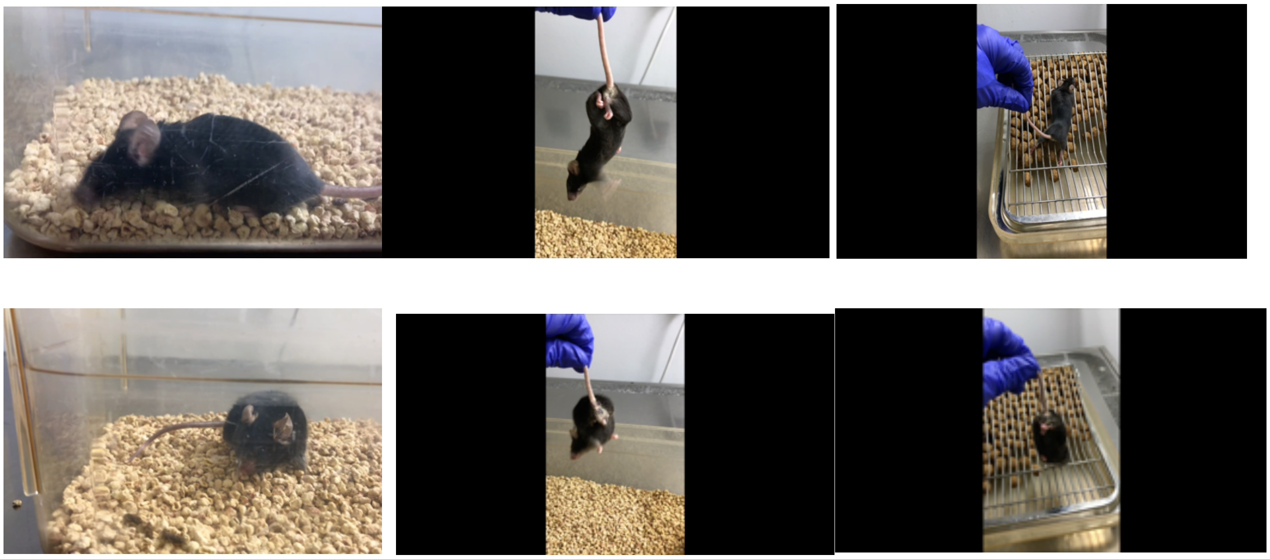

**Movie S1. Clinical performance of 4-month-old and 12-month-old control and Mettl14 cKO mice**

Representative videos demonstrating tremor and hindlimb clenching and activity levels in Mettl14 conditional knockout (cKO) mice at 4 months and 12 months of age. These videos highlight the differences in movement patterns, gait, and overall neuromuscular function across age groups.
